# GephyrinΔ199-233 - an epileptogenic microdeletion

**DOI:** 10.1101/2025.08.26.672322

**Authors:** Maria-Theresa Gehling, Laura Buchwald, Fynn Eggersmann, Filip Liebsch, Peter Kloppenburg, Günter Schwarz

## Abstract

Gephyrin, as the main organizer of inhibitory synapses, is crucial for inhibitory signal transmission, and implicated in various neurological disorders. Various studies have identified gephyrin microdeletions in conditions of autism, schizophrenia, and epilepsy. Those deletions affected the N-terminal G-domain and/or the central C-domain of gephyrin while the receptor binding C-terminal E-domain was not affected. Here, we investigated the importance of a specific microdeletion (Δ199-233) within the C-domain using a full-body knock-in mouse model. Homozygous mice displayed a severe phenotype characterized by reduced fertility, increased mortality, and neurological deficits at early developmental stages. Analyses in dissociated hippocampal neurons demonstrated disrupted synaptic targeting of gephyrin Δ199-233 that harbors the functionally important S-palmitoylation site at Cys212. Simultaneously, we found adaptations at the excitatory synapse, with smaller, but more numerous clusters of the excitatory scaffolding protein PSD95. Although, gephyrin Δ199-233 showed unexpectedly a facilitated receptor interaction, inhibitory signal transmission was reduced. We hypothesize, that the gephyrin Δ199-233-mediated reduction of inhibition triggers compensatory excitation, which possibly fails and/or disrupts the excitation/inhibition ratio in our mouse model. These findings highlight the critical role of the gephyrin C-domain and its post-translational modifications in synaptic function and neuronal health, offering a novel mouse model for the development of potential therapeutic targets addressing gephyrin-associated neurological disorders.

## Introduction

Neuronal signal transmission depends on the precise localization of receptors at the post-synaptic site. Inhibitory glycine receptors (GlyRs) and a subset of GABA type A receptors (GABA_A_Rs) are clustered by the scaffolding protein gephyrin (geph, GPHN) at the post-synapse [1]. Gephyrin is a trimeric, cytosolic protein and consists of three domains, the E-, C- and G-domain. Dimerization of the E-domain and trimerization of the G-domain allow gephyrin to self-assemble in complex multimers [2, 3]. Moreover, the dimer interface of the E-domain builds the binding pocket for the intracellular domains (ICDs) of the receptors, which are both pentameric ion channels[4]. Although the composition of receptor types varies, the interaction with gephyrin appears via the β- subunit ICD in case of GlyRs [3], and the α1-3 and β1-2 subunit ICDs in case of GABA_A_Rs [5-7]. Notably, the ICDs of both receptor types share essential structural elements required for gephyrin interaction [8, 9]. Moreover, the γ2 subunit of GABA_A_Rs was identified as important for synaptically functional receptors and the interaction towards gephyrin even though indirectly [10, 11].

Dysfunctions of gephyrin are associated with different neuronal impairments, such as epilepsy [12, 13], anxiety [14], schizophrenia [15], autism [15] and hyperekplexia [16]. Besides its function at inhibitory post-synapses, gephyrin is also expressed in peripheral tissue where it catalyzes the final steps of molybdenum cofactor (Moco) biosynthesis [17]. Defects in Moco biosynthesis led to the loss of four different Moco-dependent enzymes of which the deficiency in sulfite oxidase is the primary contributor to a severe neurodegenerative phenotype. Due to the accumulation of toxic sulfite the amino acid derivative S-sulfocysteine (SSC) [18-20] is formed causing hyperexcitation and calcium-dependent calpain-mediated cleavage a various neuronal proteins including gephyrin [20].

Besides proteolytic cleavage, clustering, localization and receptor binding of gephyrin are regulated by a large variety of post-translational modifications (PTMs), which are mostly targeting gephyrin C-domain. Among the reversible PTMs are several phosphorylation sites, involved in complex signaling cascades [21-23]. S-palmitoylation of gephyrin at cysteine 212 and 284 was found to be crucial for synaptic targeting [24]. Additionally, *cis-trans*-isomerization of a proline-rich region is catalyzed by the interaction with peptidyl-prolyl *cis-trans* isomerase 1 (Pin1) and was shown to be important for GlyR clustering [25]. Furthermore, the nucleotide exchange factor collybistin and the dynein-light chain 1/2 (DLC1/2) [26] bind to the C-domain of gephyrin, the latter may influence the transport of gephyrin and GlyRs and thus glycinergic signal transmission [27], while the importance of this interaction for GABAergic synapses is less clear. Importantly, the majority of gephyrin’s PTMs have been investigated *in vitro* [28], and only a few were linked to physiological outcomes *in vivo* [29]. Besides that, studies of human cases focused on the G-and E-domain due to their importance for receptor binding and self-assembly [12, 15, 30] while the role of the regulatory C-domain remained largely unknown. Thus, our goal was to investigate the role of the C-domain *in vivo*.

We generated a full-body knock-in mouse model causing a truncation in gephyrin C-domain spanning residues 199-233 (Δ199-233). The deleted sequence included the proposed DLC1/2 binding motif, S-palmitoylation site Cys212 and the last three residues of the *cis-trans*-isomerization motif. Characterization of homozygous GPHNΔ199-233 mice revealed that the deleted region was important for murine neuronal development and survival. We found alterations of gephyrin-receptor binding towards a tighter interaction, but reduced synaptic targeting of gephyrin due to reduced S-palmitoylation, which lead to decreased inhibitory synaptic transmission. Assumably, to overcome altered inhibition, we found adaptions at excitatory synapses and hypothesize that an altered ratio of excitation and inhibition contributes to the neuronal impairments of our mouse model.

## Results

### Severe phenotype of homozygous gephyrin Δ199-233 mice

Previously, S-palmitoylation has been identified as one important PTM to regulate synaptic plasticity of gephyrin [24]. S-palmitoylation was assigned to cysteines 212 and 284, while Cys212 being the primary site [24] (Figure 1A and B). Originally, our aim was the introduction of single amino acid substitution (C212S) allowing us to investigate solely the physiological importance of S-palmitoylation on that Cys212 residue *in vivo*. Consequently, the CRISPR/Cas9 strategy was designed to achieve the Cys212 to Ser substitution (Figure 1C). Unexpectedly, using the designed guide RNA (gRNA) and repair template in CRISPR/Cas9 system resulted in a genetic variety. In the founder generation, we identified the desired substitution C212S, but also a set of INDELs (Figure S1A). The survival rate of the founder litter was low with 25% at P20 (Figure S1C), and one mouse had to be sacrificed at P20 due to a severe seizures. Considering the high mortality, we concluded that the area we aimed to modify is important for gephyrin function. However, no mouse with a single C212S variant survived and only one mouse with a larger deletion produced offsprings, which was sufficient to establish a new line. In the second heterozygous generation we verified that the gephyrin deletion was spanning residues 199-233 (Δ199-233) and off-targets could be excluded (Figure S1C). We decided to investigate the deletion (Δ199-233) further, since several microdeletions were observed in human patients causing neuronal disorders of different kind and severity [12, 15] (Figure S1D).

**Figure 1.**
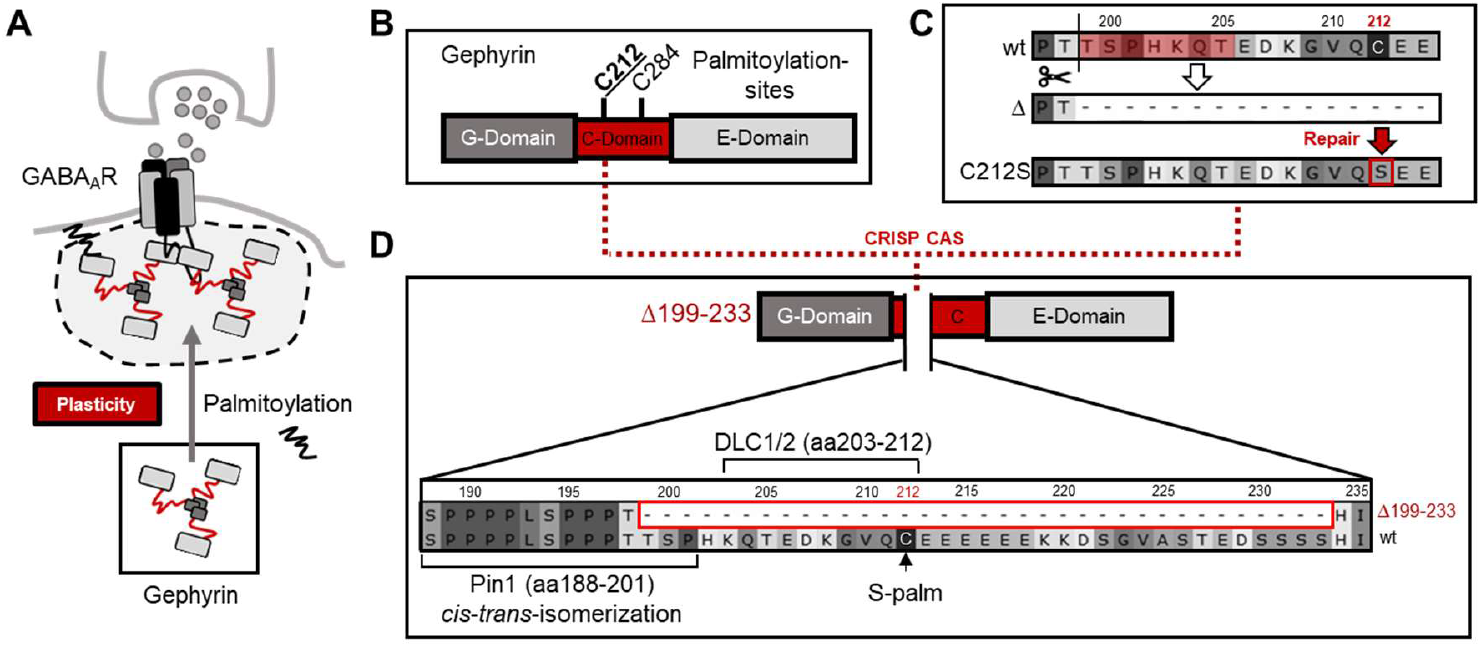
Generation of mice with microdeletion gephΔ199-233. A) Schematic description of S-palmitoylation of gephyrin influencing synaptic plasticity at GABAergic synapses. S-palmitoylation leads to gephyrin synaptic targeting and cluster size increase. B) Domain structure of a gephyrin monomer: S-palmitoylation targets are cysteines 212 and 284 in the C-domain. C212 (bold and underlined) was shown to be more relevant than C284. C) CRISPR/Cas9 strategy to obtain C212S KI mice visualized on the protein level. The potential cutting site of CAS is indicated by a scissor. Inefficient repair results in a truncated gephyrin species (Δ). Successful repair results in C212S substitution. D) Specific microdeletion of gephyrin Δ199-233 enlarged highlighting deleted PTM sites (e.g. S-palm=S-palmitoylation) and protein interaction motifs (Pin1=peptidyl-prolyl *cis-trans* isomerase 1; DLC1/2=dynein-light chain 1/2).

The truncated C-domain part is with 35 aa only a small part of the whole C-domain (residue 166-329) and includes a few PTM sites with assigned but also yet unassigned functions, which have been investigated mainly *in vitro* [24-26, 28, 31, 32]. Among the PTMs are the last three amino acids (residue 199-201) of the Pin1 interaction site, the interaction site of DLC (residue 203-212), and the S-palmitoylation site C212 (Figure 1D). Abolishing those PTMs could give new insights into their importance for synaptic plasticity *in vivo*. This would highlight the importance of regulation through gephyrin’s C-domain beside oligomerization and receptor binding abilities of G- and E-domain, respectively. Thus, we started to characterize the phenotype of Δ199-233 mice.

We found a genotype distribution of 56.3% wt/wt, 27.3% wt/Δ, but only 16.4% Δ/Δ (Figure 2A). Considering mendelian rules of crossing wt/Δ mice, we expected 25% Δ/Δ mice, thus it might be that approx. half of Δ/Δ embryos were absorbed in the uterus. Moreover, Δ/Δ pups born had a lower survival rate and often died within the first 1-4 days (Figure 2B). Following those days, 50% of Δ/Δ reached an age of P16, while 75% wt/Δ and wt/wt litter mates reached P16. In addition, approximately every third litter with at least one homozygous Δ199-233 mouse (Δ/Δ litter) completely died after P2 (Figure 2C). In comparison, in litters containing no Δ/Δ mouse (ctrl litter), approx. 90% of the pups survived P2 and no litter completely died before P2. This observation might be due to enhanced stress on maternal care induced by the sensation of impaired offspring. However, Δ/Δ mice surviving P2 did not reach weaning age P21 either. We observed that five individual mice suffered either from sudden death (P19) or had to be sacrificed in order to reduce stress levels given by their epileptic-like seizures (P17) (Figure 2D), impaired locomotion (P14-19), apathetic behavior (P18) or severe weight loss (P16). Thus, we decided to reduce animal harm and sacrificed Δ/Δ mice for experimental use before severe impairments manifested regularly. As a result, we could use only 12 Δ/Δ mice out of 164 born mice from wt/Δ × wt/Δ mating for organ harvest. In general, Δ/Δ mice were significantly lighter and thus smaller in comparison to heterozygous or wt litter mates (Figure 2E and F). Heterozygous mice were indistinguishable from wt littermates, a dominant negative effect of truncated Δ199-233 on wt gephyrin was rather unlikely and we determined heterozygous mice as wt-like.

**Figure 2.**
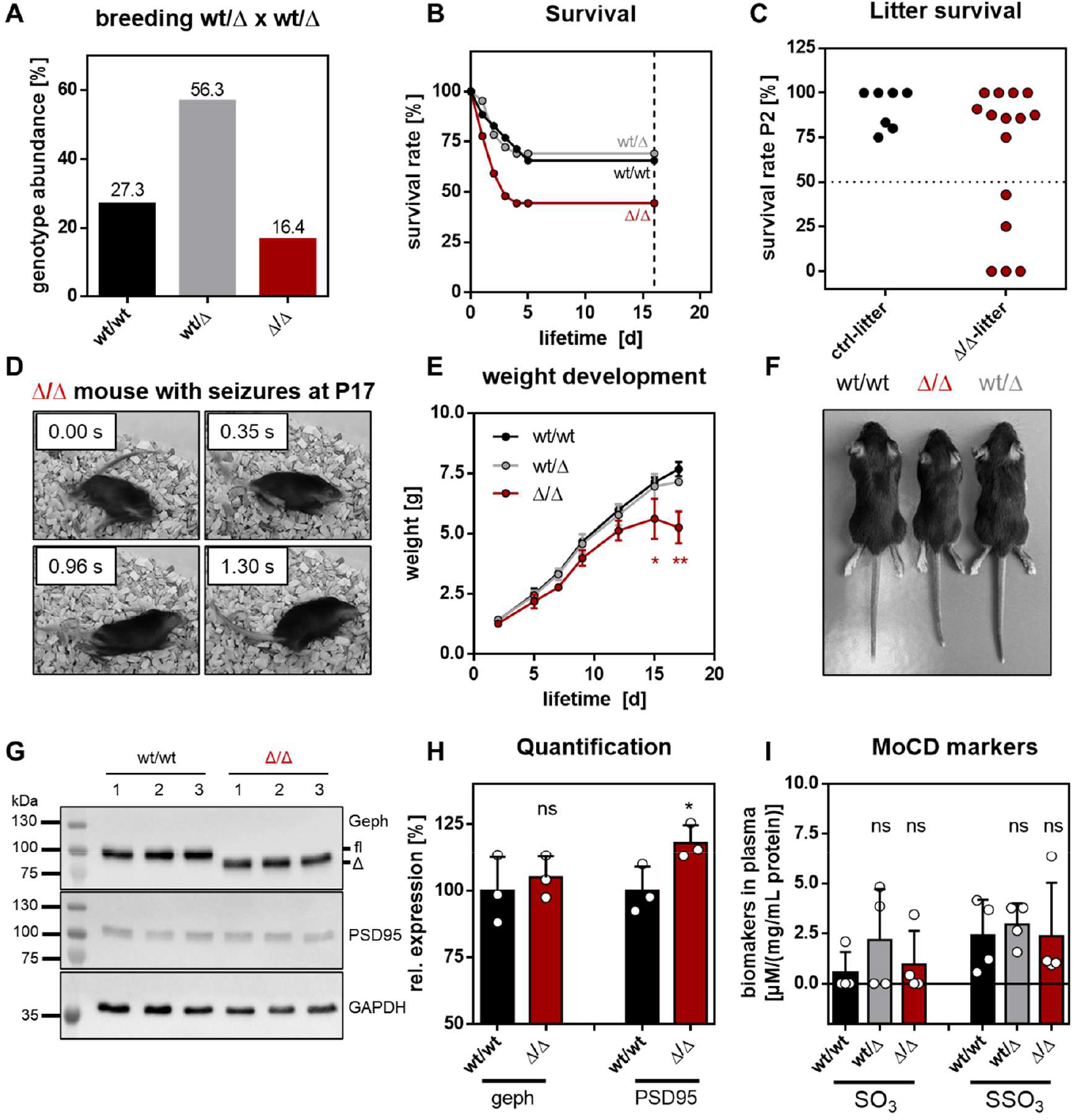
Phenotype of mice with microdeletion gephΔ199-233. A) Abundance of each genotype obtained from crossing wt/Δ × wt/Δ animals. Analysis of 164 animals. B) Survival rate of the different genotypes. Dashed line indicates analyzed animals with natural deaths and without intervention due to high stress levels (wt/wt n=35, wt/Δ n=65, Δ/Δ n=27). C) Survival rate at P2 of pups when the litter contained no Δ/Δ animal (ctrl litter, n=7) or at least one Δ/Δ animal (Δ/Δ litter, n=15). D) Images taken during a seizure episode of a homozygous GephΔ199-233 animal at P17: Rapid extension and contraction of limbs within a few milliseconds. E) Weight development of the different genotypes. Significance tested by 1way ANOVA *F*(2,14)=8.118; *p*=0.0046. *Bonferroni* post-hoc test: wt/wt vs. wt/Δ : *p>*0.9999 (ns, grey); wt/wt vs. Δ/Δ: *p*=0.0077 (**, red). Error bars determined by standard deviation. F) Litter mates of different genotypes at P17. G) Western Blot of brain lysates at P17 (n=3 per genotype). Upper panel: Gephyrin wt (fl) and truncated Δ199-233 species. Middel panel: PSD95. Lower panel: Loading ctrl GAPDH. H) Quantification of band intensities of G. Intensities of geph or PSD95 corrected by intensity of GAPDH and normalized to the average of wt/wt samples. Significance tested with *student’s t-test*: geph *p*=0.5984 (ns); PSD95 *p*=0.0492 (*). Error bars determined by standard deviation. I) Analysis of Moco deficiency (MoCD) biomarkers (SO_3_ and SSO_3_) in plasma (n=4 mice per genotype). Values normalized to protein content in plasma. Error bars determined by standard deviation.

Next, we checked protein levels of gephyrin in comparison to PSD95 in total brain lysates (Figure 2G). We found that in Δ/Δ mice the levels of gephyrin were unchanged in comparison to wt/wt mice, but the levels of PSD95 were slightly but significantly increased by approx. 20% (Figure 2H). Lastly, since weight loss, neuronal impairments and early mortality can be signs of MoCD, we tested whether Δ199-233 mice showed elevated levels of MoCD biomarkers sulfite (SO3) and thiosulfate (SSO3). The levels of these markers in blood plasma of animals at P16 (Figure 2I) were normal in all genotypes. Thus, we excluded a contribution of MoCD to this phenotype.

In summary, the phenotype of Δ/Δ mice was lethal and appeared at early developmental stages. Mostly, at an age of P16-P18 mice showed different signs of severe impairments (seizures, sudden death), most obvious a reduced body weight. The reason for sudden death and weight loss was not obvious, but we assumed that neuronal dysfunctions were the cause.

#### Altered post-synaptic scaffolds in gephyrin Δ199-233-expressing neurons

Breeding success to gain homozygous gephyrin Δ199-233 mice to analyze gephyrin clustering was limited, so we decided to overexpress recombinant mScarlet-tagged gephyrin wt and Δ199-233 in hippocampal cell cultures derived from conditional gephyrin knockout (GephyrinFlox) mice. We used adeno-associated viral transduction for efficient expression of the transgene. Identification of GABAergic synapses was performed by staining of vGAT and GABA_A_R γ2 (Figure 3A). For excitatory synapses we used PSD95 and vGLUT (Figure 4A). Via automated image analysis [33], scaffold cluster sizes, intensities and number of synaptic and non-synaptic punctae were determined (Figure 3B-G and Figure 4B-E).

**Figure 3.**
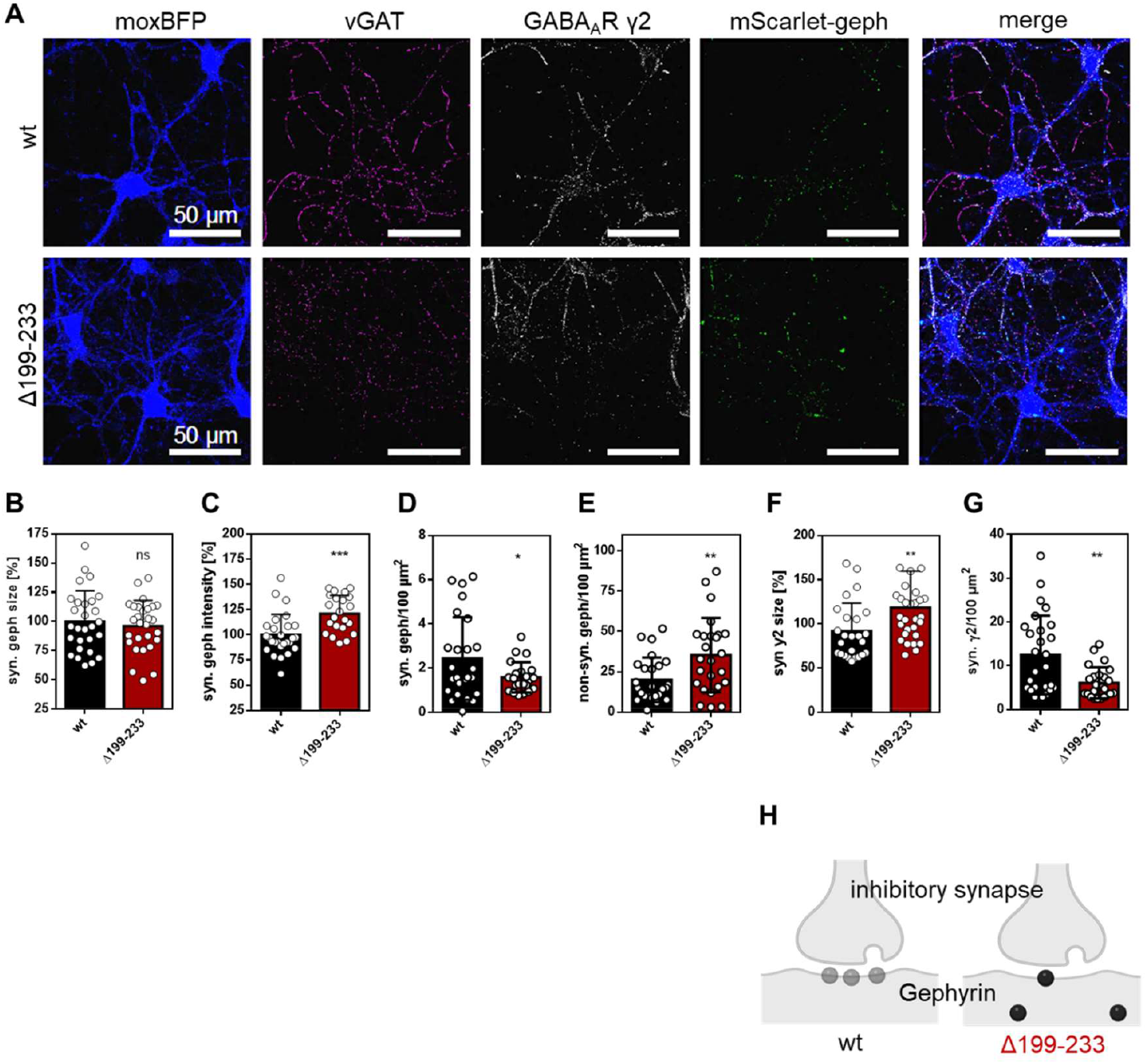
Inhibitory synapses in gephΔ199-233-expressing neurons. Expression of mScltGeph wt and Δ199-233 in primary hippocampal neurons of GephFlox mice. Endogenous Geph KO induced by moxBFP-P2A-Cre. Images were processed with equal brightness thresholds and DOGs in all channels except of moxBFP. Cluster analysis: Significance tested by *student’s t-test*. Error bars determined by standard deviation. Analysis of inhibitory GABAergic synapses. Analysis of n=30 cells per condition. A) Images of representative neurons. Synaptic GABAergic gephyrin identified by co-localization with vGAT and GABA_A_R γ2. B) Sizes of synaptic gephyrin punctae normalized to average size of wt. *p*=0.3136 (ns). C) Intensity of synaptic gephyrin punctae normalized to average intensity of wt. *p*=0.0004 (***). D) Number of synaptic gephyrin punctae per 100 μm^2^ cell area. *p*=0.0488 (*). E) Number of non-synaptic gephyrin punctae per 100 μm^2^ cell area. *p*=0.0066 (**). F) Sizes of synaptic GABA_A_R γ2 punctae normalized to average size in wt condition. *p*=0.0099 (**). G) Number of synaptic GABA_A_R γ2 punctae per 100 μm^2^ cell area. *p*=0.0022 (**). H) Schematic illustration of synaptic clustering Δ199-233 gephyrin in comparison to wt: Cluster sizes are maintained, but intensity increases, less synaptic clusters occur, while non-synaptic clusters accumulate.

**Figure 4.**
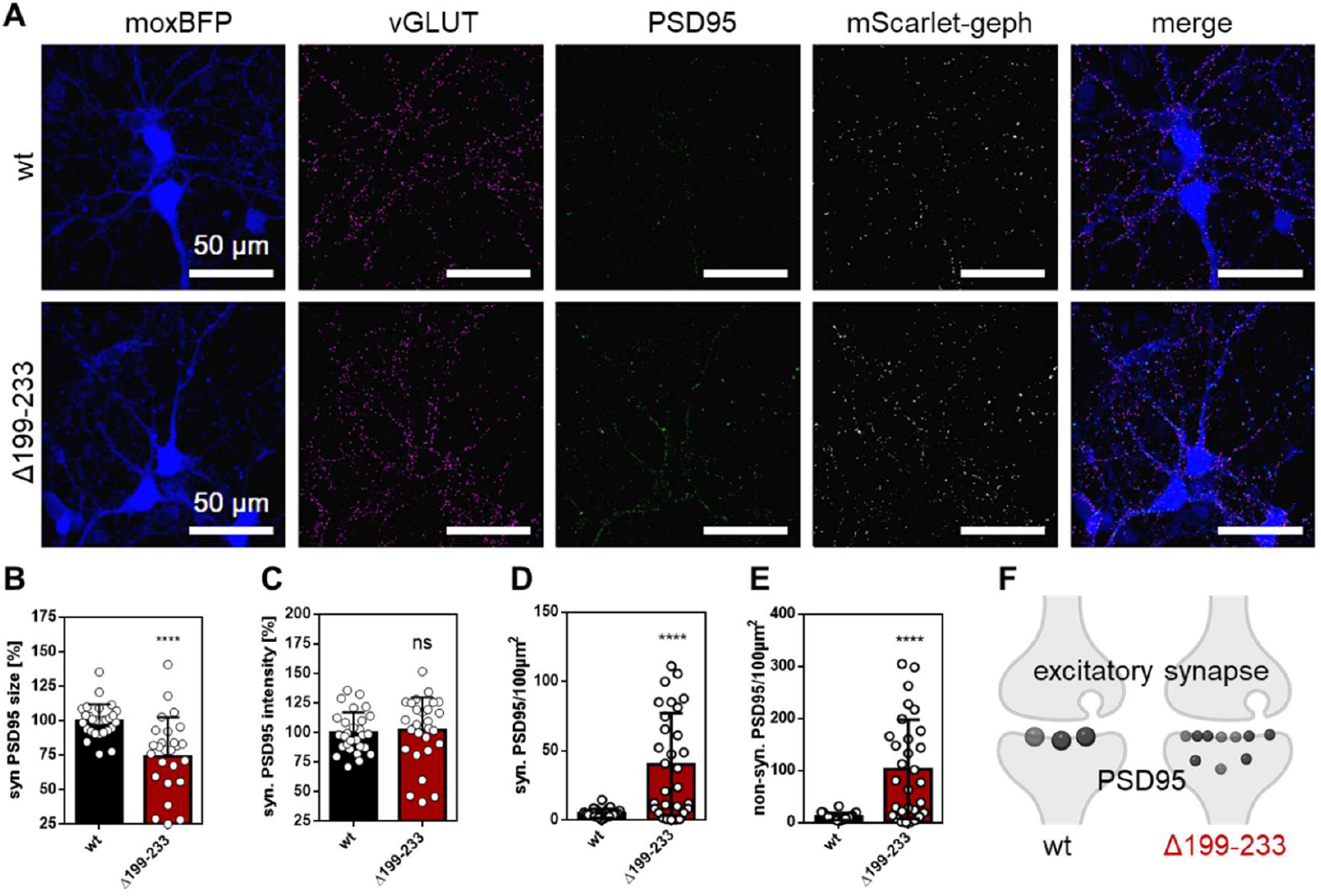
Excitatory synapses in gephΔ199-233-expressing neurons. Expression of mScltGeph wt and Δ199-233 in primary hippocampal neurons of GephFlox mice. Endogenous Geph KO induced by moxBFP-P2A-Cre. Images were processed with equal brightness thresholds and DOGs in all channels except of moxBFP. Cluster analysis: Significance tested by *student’s t-test*. Error bars determined by standard deviation. Analysis of excitatory synapses. Analysis of n=30 cells per condition. A) Images of representative neurons. Synaptic PSD95 punctae identified by co-localization with vGLUT. B) Sizes of synaptic PSD95 punctae normalized to average size of wt. *p<*0.0001 (****). C) Intensity of synaptic PSD95 punctae normalized to average intensity of wt. *p*=0.7221 (ns). D) Number of synaptic PSD95 punctae per 100 μm^2^ cell area. *p<*0.0001 (****). E) Number of non-synaptic PSD95 punctae per 100 μm^2^ cell area. *p<*0.0001 (****). F) Schematic illustration of synaptic clustering of PSD95 in Δ199-233 gephyrin expressing neurons in comparison to wt expressing cells: Cluster intensities are maintained, but sizes decreased, more synaptic and non-synaptic clusters occur.

At GABAergic synapses, we observed that the average fluorescence of gephyrin Δ199-233 punctae was significantly more intense in comparison to gephyrin wt (Figure 3C), but the sizes of punctae stayed comparable (Figure 3B)). The number of non-synaptic gephyrin Δ199-233 punctae increased significantly (Figure 3E), while the number of corresponding synaptic punctae significantly decreased (Figure 3D). This indicated that synaptic targeting of gephyrin Δ199-233 is likely disturbed, which fits to the lack of S-palmitoylation at C212 in gephyrin Δ199-233 [24]. Furthermore, we observed that the sizes of GABA_A_R γ2 punctae at synapses increased significantly (Figure 3F), but significantly less synaptic GABA_A_R γ2 punctae could be found per area. This indicated that the interaction of receptor and gephyrin could be tighter, while synaptic targeting was disturbed, which could affect synaptic transmission in summary.

At excitatory synapses, we observed in gephyrin Δ199-233 expressing cells that PSD95 punctae were significantly smaller (Figure 4B) than those of gephyrin wt expressing cells. The average intensity stayed comparable (Figure 4C). Remarkably, the number of synaptic as well as of non-synaptic PSD95 punctae significantly increased (Figure 4D and E) in gephyrin Δ199-233 expressing neurons. These findings suggested that PSD95 scaffolds were split in smaller fractions, leading to overall more punctae (Figure 4F). Assumably, neurons aim to “balance” excitation and inhibition, which might be triggered though the observed changes at the inhibitory synapse.

To investigate the effects on inhibitory transmission, we performed electrophysiology experiments with the same experimental set up of neurons. We observed a tendency of alterations of miniature inhibitory post-synaptic currents (mIPSCs) (Figure 5). Surprisingly, the frequency of mIPSCs in gephyrinΔ199-233 expressing cells was comparable to gephyrin wt expressing neurons indicating an unchanged number of receptors present at the post-synapse. First, this seems contra-intuitive to the previous imaging analysis of reduced numbers of synaptic GABA_A_R γ2 punctae. But considering the increase of sizes of synaptic GABA_A_R γ2 punctae, it is possible, that still the same “amount” of GABA_A_Rs was present to generate mIPSCs. Consequently, both parameters, number and size, are decisive for inhibitory signal transmission. Additionally, we observed a slight decrease of amplitude (Figure 5E), but surprisingly an increase of decay time (Figure 5D). The integration of mIPSCs displays the presence of negative charges at the post-synapse, which was decreased for gephyrinΔ199-233 expressing cells (Figure 5C). Assumably, the gephyrinΔ199-233- receptor-interaction might be changed as previously assumed resulting in decreased inhibitory neurotransmission.

**Figure 5.**
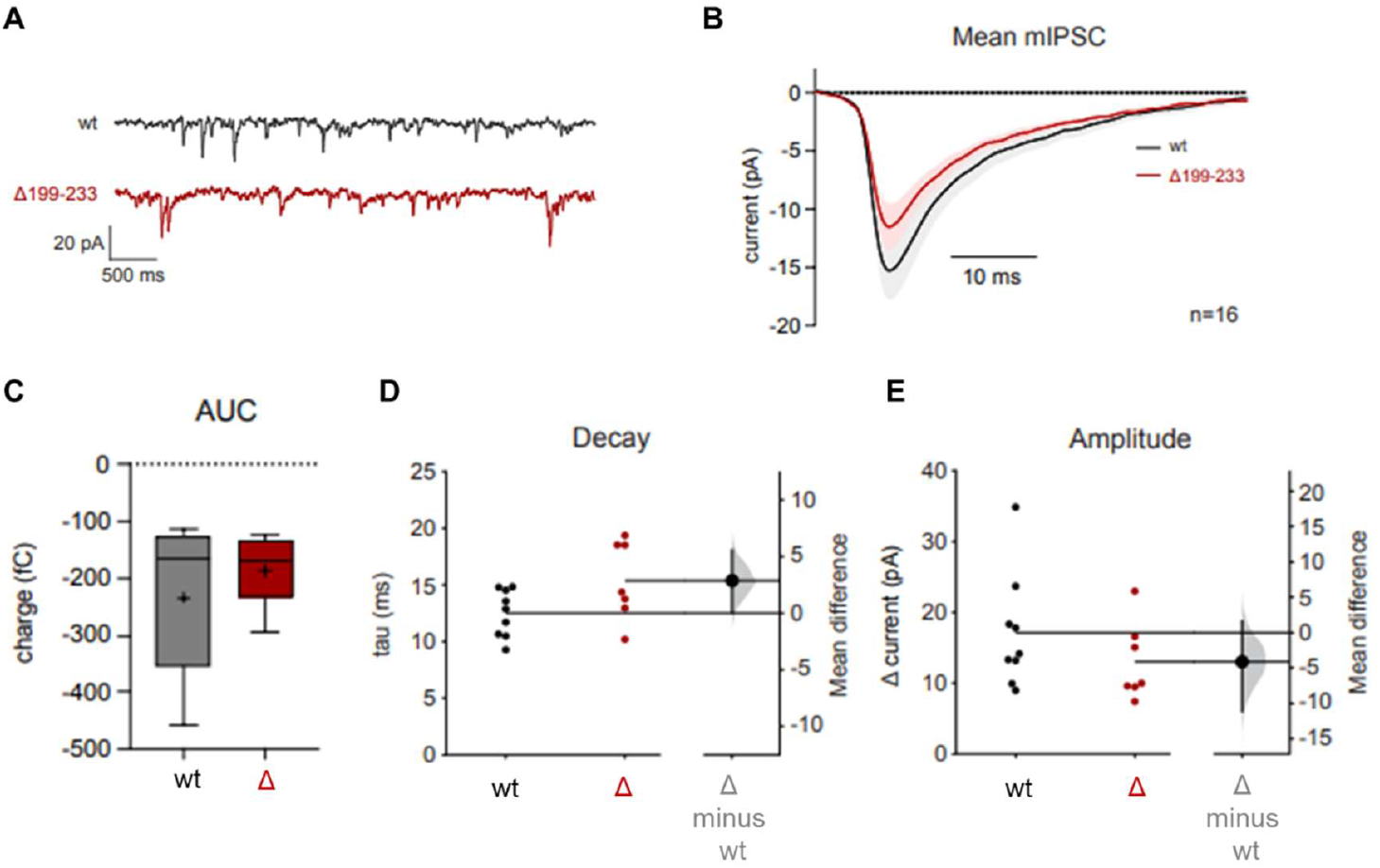
Inhibitory signaltransmission in gephΔ199-233-expressing neurons. Measurements and analysis of miniature inhibitory post-synaptic currents (mIPSCs) in in primary hippocampal neurons of GephFlox mice. Expression of mScltGeph wt and Δ199-233. Endogenous Geph KO induced by moxBFP-P2A-Cre. Analysis pf 16 cells per condition. Statistical analysis by mean difference of the confidence interval and *p*-value permutation test. Error bars determined by standard deviation. A) Representative mIPSCs of wt and Δ199-233 cells. B) Average mIPSCs. C) Quantification of the area under the mIPSC curve (AUC). D) Quantification of mIPSC decay time. E) Quantification of mIPSC amplitude.

In summary, both types of synapses were affected in gephyrin Δ199-233 expressing neurons. On the one hand, GABAergic synapses contained less synaptically localized but bigger gephyrin and GABA_A_R γ2 punctae combined resulting in reduced inhibitory signal transmission. On the other hand, excitatory synapses contained more synaptically localized, but smaller PSD95 punctae. These changes at both synapses might resemble a disturbance of E/I ratio in Δ/Δ mice, which is a possible explanation for a neuronal impaired phenotype.

### Disturbed S-palmitoylation and DLC interaction of gephyrin Δ199-233

Since several PTMs are affected by the microdeletion in gephyrin Δ199-233, we investigated the contribution of distinct PTMs. First, we co-expressed EGFP-gephyrin and mCherry-DLC1 in COS7 cells (Figure 6) according to the study, when the DLC binding region of gephyrin was originally identified [26]. DLC is recruited into gephyrin clusters when gephyrin wt was expressed. In contrast co-expression with gephyrin Δ199-233 led to a diffusely distribution of DLC (Figure 6A). Quantification of the co-localization with Mander’s co-localization coefficient (MCC) resulted in significantly less gephyrin Δ199-233 overlapping DLC (M1) and DLC overlapping gephyrin Δ199-233 (M2) in comparison to gephyrin wt (Figure 6B). The low affinity binding site of gephyrin for DLC is left in gephyrin Δ199-233, but this is not sufficient to enable an interaction. This is line with the first study of the gephyrin-DLC interaction, when only the high-affinity binding site was missing resulting in no interaction in several experiments [26]. Here, we verified that the binding of gephyrin Δ199-233 and DLC1 is disturbed due to the lack of the high affinity binding motif.

**Figure 6.**
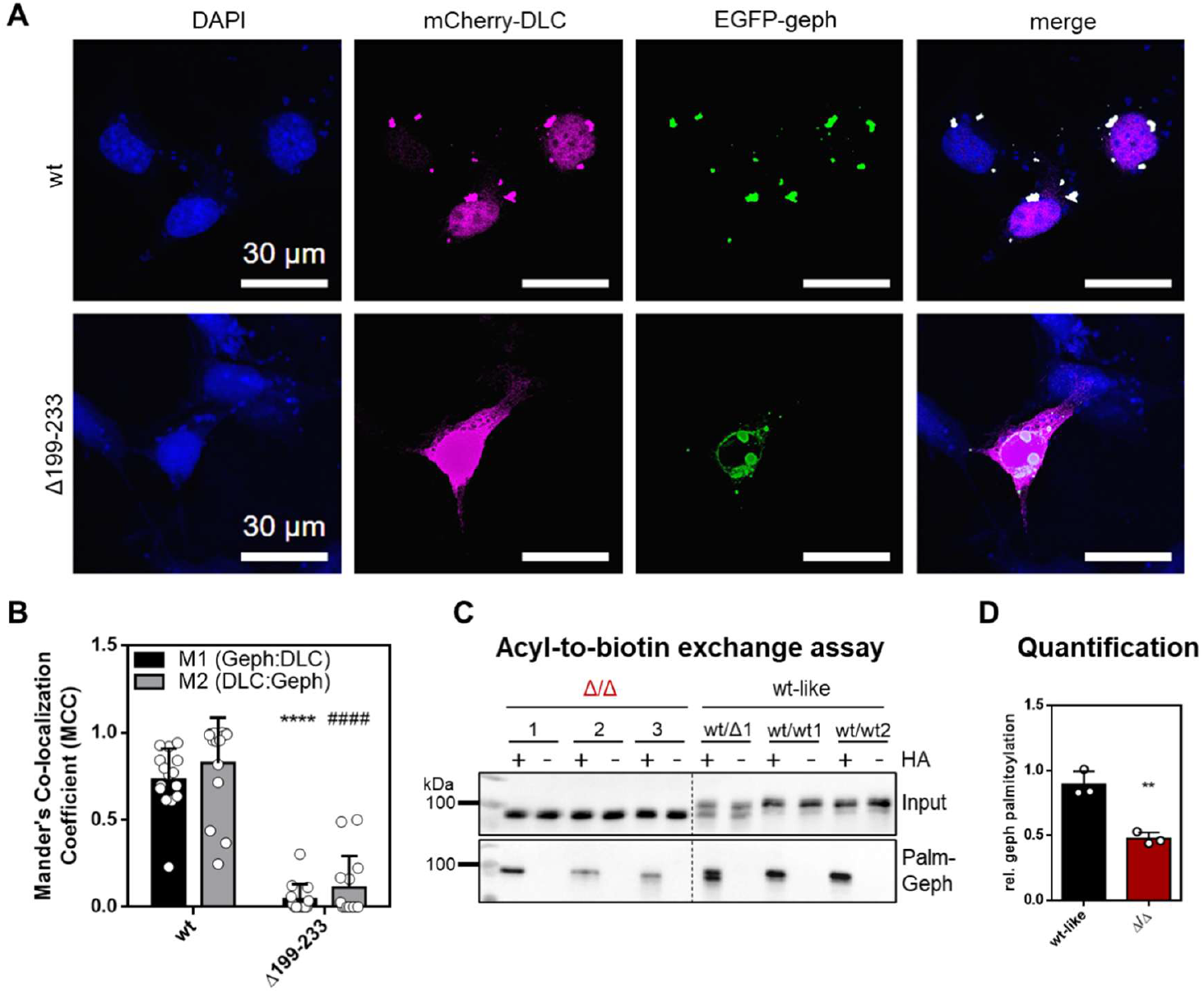
Characterization of selected PTMs and interaction partners of gephΔ199-233. A) Analysis of gephyrin S-palmitoylation via acyl-to-biotin exchange assay. S-palmitoylated gephyrin species (lower panel; Palm-Geph). Analysis of brain lysates (n=3 mice per condition). B) Quantification of band intensities of A. Palm-Geph normalized to input signal. Significance tested by *student’s t-test p*=0.0030 (**). Error bars determined by standard deviation. C) Pull-down of endogenous gephyrin from brain lysates. Analysis of co-pulled binding partners. D) Representative images of COS7 cells expressing mCherry-DLC1 and EGFP-gephyrin wt and Δ199-233 respectively. Nuclei stained with DAPI. E) Quantification of co-localization of gephyrin and DLC using Mander’s co-localization coefficient (MCC). M1: Gephyrin overlapping DLC; M2: DLC overlapping gephyrin. Significance tested by *student’s t-test*, significance level alpha corrected by the number of tests (2). M1 wt vs M1 199-233: *p*<0.00005 (****); M2 wt vs M2 199-233: *p*<0.00005 (####). Analysis of n=15 cells per condition.

Furthermore, we analyzed the S-palmitoylation levels of gephyrin in full brain lysates of Δ/Δ mice in comparison to wt/wt and wt-like mice (Figure 6C and D). The biotin-to-acyl exchange assay revealed, that the S-palmitoylation level of gephyrin Δ199-233 was significantly reduced to half of that of gephyrin wt (Figure 6D). With this we verified our expectation that the impairment of synaptic trafficking observed in previous *in cellulo* experiments could be explained by the lack of S-palmitoylation [24].

### Gephyrin Δ199-233 shows facilitated receptor interaction

Next, we characterized the microdeletion *in vitro* using recombinantly expressed gephyrin from E. coli. First, we examined secondary structure by circular dichroism spectroscopy (Figure 7A). We found the dominance of α-helices with minima around 210 and 220 nm for both gephΔ199-233 and wt. Second, oligomerization was investigated by size exclusion chromatography (Figure 7B and C). As expected, Gephyrin Δ199-233 eluted later than gephyrin wt, due to the deletion of 34 aa per monomer. Since, gephyrin appears at minimum in a trimeric state (Figure 7B) when derived from E.coli [28], the deletion adds up to 102 aa, which mirrors a difference in the hydrodynamic radius of approx. 80 kDa. The more complex and higher the oligomeric state becomes, the bigger gets the apparent weight difference between gephyrin Δ199-233 and wt (Figure 7C). However, gephyrin Δ199-233 was still able to self-assemble and form networks.

**Figure 7.**
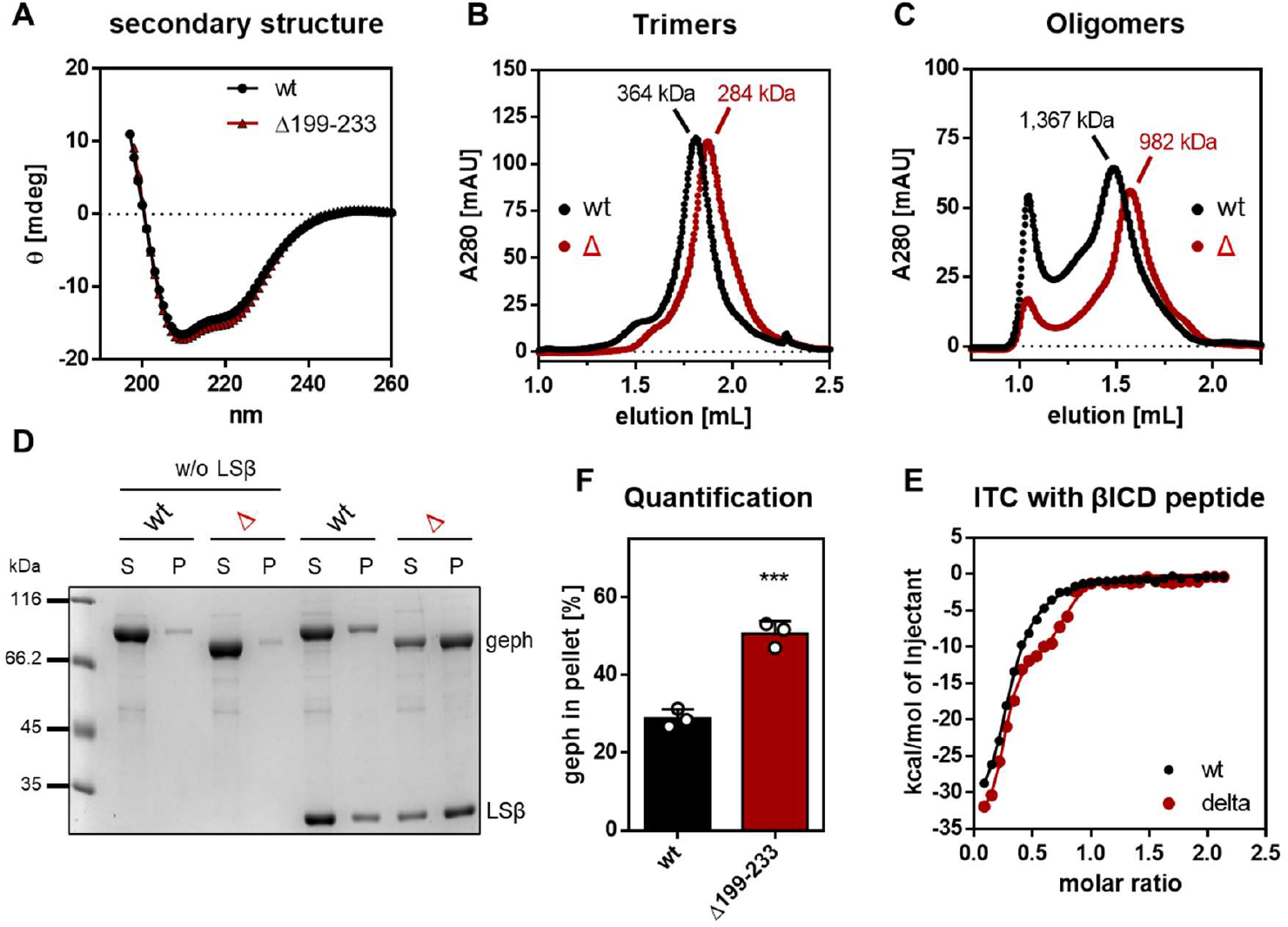
Characterization of GephΔ199-233 *in vitro*. Experiments performed with recombinantly expressed gephyrin wt and Δ199-233 from *E*.*coli*. Representative data from three individual experiments using proteins from three independent purifications (n=3). A) Secondary structure measured by circular dichroism spectroscopy. B-C) Oligomeric state detected by analytical size exclusion chromatography (SEC). B) Trimers of gephyrin eluted around 1.8 mL. Apparent weight determined to 364 kDa for wt gephyrin and 284 kDa for Δ199-233. C) Oligomers of gephyrin eluted around 1.6 mL and high-order oligomers at 1 mL. Apparent weight determined to 1 367 kDa for wt gephyrin and 982 kDa for Δ199-233. D) Sedimentation assay using LS βICD as a pentameric GlyR model. E) Quantification of F. Sedimented gephyrin in pellet determined and normalized to total protein. Significance tested by *student’s* t-test: *p*=0.0007 (***). F) Isothermal calorimetry titration experiment using GlyRβICD peptide as a model for the GlyR. The peptide was titrated into gephyrin. Exemplary overlay of fitted curves of wt and Δ199-233 gephyrin. Find full plots in supplementary figure S2.

Lastly, we tested the gephyrin-receptor interaction in binding studies and observed, that this interaction seems to be facilitated for gephyrin Δ199-233. This was revealed by two different approaches: a) ITC experiments using the established GlyR β-subunit peptide [3] (Figure 7E) and b) in a sedimentation assay (adapted after Bai et al [34]) in which we used a fusion protein of the lumazine synthase (LS) and five GlyR β-subunit ICDs (LSβ) as a pentameric GlyR model [35] (Figure 5D and F). In ITC experiments gephyrin shows a two-sided interaction with the peptide resulting in a low and a high affinity binding site [3]. The dissociation constants for wt and Δ199-233 gephyrin were comparable for both binding sites (*K*_D_^high^ : wt=0.063±0.017 μm; Δ199-233=0.050±0.023 μm, *K*_D_^low^ : wt=1.83±0.059 μm; Δ199 233=1.54±0.7 μm). But the shape of the ITC run reveals a more prominent low affinity binding site (Figure 5E, Figure S2). Interestingly, when exposing gephyrin to the pentameric GlyR model LSβ in the sedimentation assay (Figure 5D), we found twice as much gephΔ199-233 in the pelleted fraction compared to gephyrin wt (Figure 5E). The interaction towards the multimeric receptor is responsible for gephyrin sedimentation [34], which indicates that this interaction is facilitated for gephΔ199-233. Although, we used model systems for the GlyR, the required structural elements of both, gephyrin-GlyR β and gephyrin-GABA_A_R α3 interaction for instance, are comparable [8]. Additionally, gephyrin does not directly bind to GABA_A_R γ2 subunits and requires additional α1-3 subunits. Thus, the results of the binding studies can be used as an explanation for the cluster analysis of expressed gephΔ199-233 at GABAergic synapses in neurons previously: Due to the facilitated gephyrin-receptor interaction, the intensity of gephΔ199-233 co-localizing with vGAT and GABA_A_R γ2 punctae was increased, while simultaneously the sizes of synaptic GABA_A_R γ2 punctae were increased.

## Discussion

We identified gephyrin with the microdeletion Δ199-233 as a new pathogenic variant in mice. This pathogenicity is evident through reduced fertility and increased mortality, characterized by weight loss, sudden death, and epileptic-like seizures. The phenotype emerged in the early developmental stage (P16-P19) and was not related to MoCD. In comparison, human patients with larger exonic microdeletions in gephyrin suffer from complex neuronal disorders such as autism, schizophrenia, and seizures, which typically manifest in early childhood [12, 15]. Our mouse model thus appears suitable for investigating similar conditions in humans and might be valuable for developing treatment strategies. However, human cases often involve deletions in the G- and E-domains of gephyrin, [12, 15] which are critical for clustering and receptor binding [2]. These domains are believed to be major contributors to these disorders, whereas the C-domain has received less attention.

Our study highlights the crucial role of the C-domain of gephyrin, which includes several PTM sites [24, 28, 36], in the onset of neuronal diseases. We investigated these PTMs to understand their physiological relevance and their contribution to the phenotype in our mouse model. The absence of PTMs in gephyrin Δ199-233 could be attributed to the lack of S-palmitoylation and DLC interaction. The significance of missing phosphorylation (T199, T205, S222, S226, and T227) remains unclear as their roles in synaptic functions of gephyrin have not been fully defined [28]. However, phosphorylation site S200 is crucial for gephyrin-Pin1 interaction. A substitution significantly reduces this interaction, although some binding activity remains [25]. *Cis-trans* isomerization of gephyrin, facilitated by Pin1, is important for recruiting synaptic GlyRs [25], which mediate inhibition in the brain stem and spinal cord [37]. This impairment would typically manifest as stiffness [38, 39], contrary to the seizures observed in our model. Therefore, we hypothesize that a residual gephyrin-Pin1 interaction in gephyrin Δ199-233, is likely maintained and does not contribute to the phenotype.

In contrast, the interaction with DLC was completely abolished in Δ/Δ mice, as demonstrated *in cellulo*. This aligns with previous studies, where gephyrin variants lacking the high affinity DLC binding motif (residue 203-212) fail to interact with DLC [26]. Interestingly, it was previously stated that the ‘DLC binding is not essential for synaptic localization of gephyrin’ although the S-palmitoylation site Cys212 was also absent in those studies, possibly affecting the results. The absence of Cys212 alone reduces synaptic gephyrin due to lack of S-palmitoylation [24]. Thus, the combined absence of both PTMs may balance each other out through an unknown mechanism.

Furthermore, a new role of DLC-gephyrin binding was recently proposed with regard to liquid-liquid-phase separation (LLPS). LLPS might contribute to the complexity of synaptic structures by the formation of microdomains [34, 40]. It was shown, that gephyrin undergoes LLPS, proposed as a key mechanism for regulating synaptic clustering [34, 40, 41]. The interaction with DLC further strengthened LLPS formation of gephyrin [34]. Thus, while the synaptic function of gephyrin might not be heavily dependent on DLC-binding, the disrupted DLC-gephyrin interaction in Δ199-233 is likely not the primary cause of neuronal impairments observed in Δ/Δ mice.

The strongest contribution of PTMs was found for S-palmitoylation. S-palmitoylation levels of gephyrin Δ199-233 were significantly reduced to 50% *in vivo*, due to the missing residue Cys212. The second S-palmitoylation site Cys284 was unable to fully compensate, which is in line with previous reports [24]. Lack of S-palmitoylation resulted previously in reduced synaptic targeting and cluster sizes of gephyrin, impairing GABAergic neurotransmission [24]. Although gephyrin Δ199-233 showed reduced synaptic localization, cluster sizes were not reduced, suggesting an interplay of opposing PTM effects or another mechanism. *In vitro* binding studies indicated that the gephyrin-receptor interaction via the low affinity binding site was facilitated for gephyrin Δ199-233, possibly because the shorter C-domain no longer masked this site. This hypothesis is supported by previous studies, which suggested a direct modulation of the C-domain for gephyrin-receptor interaction independent on further PTMs [3, 42].

In conclusion, while gephyrin Δ199-233 was impaired in synaptic recruited likely due to reduced S-palmitoylation, a robust proportion still reached the synapse and interacted more strongly with receptors. Still, this proportion was not sufficient to maintain normal inhibition, since we observed reduced GABAergic transmission in electrophysiology experiments. This suggests that PTM-driven synaptic targeting, such as S-palmitoylation, may be crucial for gephyrin’s function over mere receptor interaction.

However, this hypothesis would lead to a reduction of the frequency of mIPSCs, which we did not observe. Instead, we observed a longer decay time. Consequently, we postulate that enough GABA_A_Rs are clustered at the synapse, but their kinetics might be altered. Previous studies support this hypothesis, since it was suggested, that the receptor ICD contacts gephyrin C-domain [3]. Thus, it is possible, that the flexible C-domain not only masks the low-affinity receptor binding site, but also interferes with the chloride influx of GABA_A_Rs. Regardless of the underlying molecular reason, a reduced GABAergic transmission alone is only one explanation for the observed phenotype in Δ/Δmice.

To further understand the phenotype of Δ/Δ mice, we considered the interplay of excitation and inhibition (E/I) and examined excitatory synapses. We observed equal expression of truncated and wt gephyrin *in vivo*, but slightly increased levels of the excitatory scaffolding protein PSD95 suggesting an alteration of E/I. Increased synaptic and non-synaptic PSD95 punctae with reduced sizes were also noted in cell culture experiments, supporting potential adaptations in E/I [43, 44]. As a result, complex neuronal impairments due to morphological changes of neuronal architectures could appear in Δ/Δ mice [45, 46], especially when the adaptions on one site cannot be mediated accordingly. Interestingly, the phenotype became visible after approx. 14 days, which is the time window when mice develop full sensory receptivity and show increased sensitivity to extrinsic provocation [47]. In this period, the functional interplay of excitation and inhibition is even more challenged [48].

In summary, we propose that the phenotype observed in Δ/Δ was caused by a reduction of inhibition due to disturbed PTM-driven synaptic targeting, which leads to adaptations at excitatory synapses (Figure 8). This adaption might work till a certain point, until it became insufficient and caused hyperexcitation, visible as epileptic-like seizures for instance. However, the exact cellular changes in further neuronal networks *in vivo*, the disturbed E/I ratio and its adaption mechanisms in Δ/Δ mice are still under our research.

**Figure 8.**
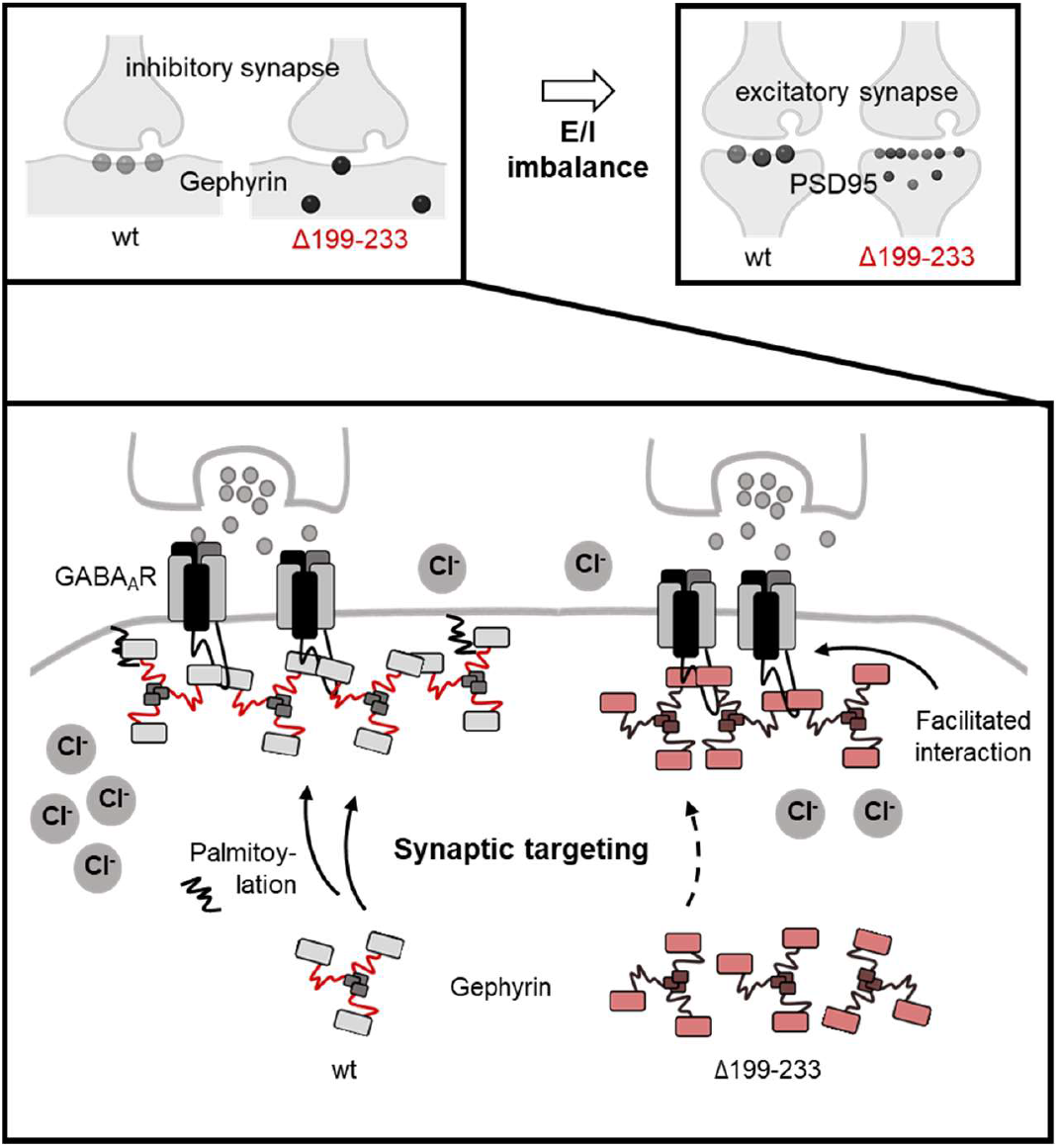
Hypothesized E/I imbalance in gephyrin Δ199-233 expressing neuronal networks. Schematic illustration of the phenotype of homozygous gephyrin Δ199-233 mice caused by an imbalance of excitation and inhibition (E/I): At GABAergic inhibitory synapses, gephyrin Δ199-233 leads to a reduced signal transmission (Cl^-^ influx). Although, the gephyrin-receptor interaction is facilitated, the recruitment by S-palmitoylation is disturbed for gephyrin Δ199-233 (indicated by a dashed arrow) leading to an accumulation of non-synaptic gephyrin. At the excitatory synapses, PSD95 shows smaller but more numerous clusters, likely an adaptation to the changes at the inhibitory synapse.

## Material and Methods

### CRISPR CAS

One step generation by electroporation in zygotes for C212S in the Gphn gene was used (DOI 10.3390/cells9051088; DOI 10.1038/s41598-017-08496-8). Used components: gRNA_1: TTATGTGGGCTAGTTGTAGG (AGG); gRNA2: TGCTGCAGTTATATGTGATG (AGG); ssODN repair template (Ultramer): A*A*GATTTACCTTCC CCACCACCTCCTCTCTCTCCACCTCCcACtACTAGCCCACATAAACAGACAGAAGACAAAGGAGTTCAGTccGAgGAAGGGAAGAAGAAAAGAAAGACAGTGGTGTAGCTTCAACAGAAGATAGTTCaTCATCACATATAACTGCAGCAGCTCTTGCTGCAAAGGTAAG T*C*T (* = phosphorothioate bonds to increase nuclease resistance).

### Genotyping and sequencing

DNA was isolated from ear punches or tail cuts using QuickExtractTM (Lucigen). The F0 generation was typed by using PCR (KAPA2G Fast Genotyping Mix (KAPA Biosystems)) (fw_TCAGTTCATACTGCCAGCTCTAC; rv_AGGCAATGACACATCATTAGTGC) followed by BsaJI digestion. The next generations were screened by PCR using fw_CCTCA TGCCATTGACCTTTTACG and rv_ATCATTAGTGCCTCAGAGCTCAA. Off-targets were screened by PCR using Chr1_fw_TGTCACGACAAAAGACAAAGATCA, Chr1_rv_CCCCACTTCTCTGGTCCCTA, Chr12_fw_CCAGATGGACCACACTAATG, Chr12_rv_TCTCCTAGGCCTACAAGACAA, Chr15_fw_AAGAAGCCTCCTTCCTCA CAAG and Chr15_rv_TGCTACATCAGACATGCCTTCTC. After agarose gel electrophoresis amplicons were extracted by the E.Z.N.A. Gel extraction Kit and sent for sanger sequencing (Eurofins).

### Sacrifice of mice and sample preparation

Mice were anesthetized with Ketamine/Xylazine through intra-peritoneal injection until decapitation could be safely done. Blood samples were collected by a Pasteur pipette, flashed with 100 mM EDTA (pH 7.5) and centrifuged for 10 min at 17 000 × g and 4°C. The supernatant (blood serum) was collected, snap frozen with liquid nitrogen and stored at -80°C.

The brain was removed and used for lysates. Therefore, RIPA buffer (50 mM Tris/HCl pH 8.0, 150 mM NaCl, 0.1% SDS, 1% IGEPAL CA-630, 0.5% Na deoxycholate, 1 mM EDTA, freshly added protease inhibitor (Roche)) was added. The tissue was dounce homogenized with 1 200 rpm 20 times, sonicated 10 sec at 4°C and 30% amplitude and subsequently centrifuged for 10 min at 4°C and 4 000 × g. The supernatant was transferred into a new tube, snap frozen with liquid nitrogen and stored at -80°C. Protein concentration was determined using BCA. Typically, 100 µg protein were used for SDS-PAGE for WB analysis, 2 mg for acyl-to-biotin exchange assays and 2 mg for pull-down assays.

### Ethics statement

We complied with all relevant ethical regulations for animal testing and research. Experiments were approved by the local research ethics committees (Germany, Landesamt für Natur, Umwelt und Verbraucherschutz Nordrhein-Westfalen, reference 2021.A089, 2019.A077 and 2021.A450).

### Determination of MoCD markers in blood serum

Blood serum samples were thaw and cleared for 10 min at 17 000 × g and 4°C. 30 μL of the supernatant were supplemented with 30 µL 100% methanol and centrifuged at 17 000 × g for 15 min at 4°C. 15 µL of the supernatant were mixed with 15 µL reaction buffer (160 mM HEPES pH 8.0, 16 mM EDTA), 15 µL 100% acetonitrile, and 3 µL 46 mM mBBr in 10% acetonitrile. The reaction was incubated for 30 min at rt (room temperature) in the dark. 30 µL methanesulfonic acid were added and the sample diluted with 312 µL running buffer A (0.25% acetic acid, pH 4). The sample was centrifuged at 17 000 × g for 15 min at 4°C and 200 µL supernatant were used for HPCL analysis. Fluorescence was measured at 380 nm, emission at 480 nm. The run was performed following the protocol in the scheme below, using a 250 × 3 mm Nucleodur HTec C18 RP column, 5 µm particle size, buffer A and B (100% methanol), an injection volume of 10 µL and a column temperature of 40°C.

### Acyl-to-biotin exchange assay

The previously described assay^23^ was changed as followed: Acetone precipitation was used to remove chemicals; 2 mg brain lysate were used and the sample after blocking (reaction with methyl methanethiosulfonate) divided in two equal samples (-HA and + HA); 100 mM hydroxylamine (HA) were used.

### cDNA Synthesis

100 mg mouse brain tissue were homogenized in 1 mL TRI Reagent (Sigma). After 5 min at rt, 100 μL 1-Brom-3-chlorpropane were added. The reaction was vortexed for 15 sec, incubated 10 min at rt, and centrifuged for 15 min at 12 000 × g and 4°C. 500 μL 2-Propanol were added to the aqueous top layer in a separate tube. The mixture was vortexed, incubated 5 min at rt and centrifuged for 10 min at 12 000 × g and 4°C. The RNA pellet was washed with 1 mL 75% ethanol and subsequently centrifuged at 7 500 × g for 5 min at 4°C. The pellet was dried at rt and resolved in 100 µL RNAse-free water. Typically, 5 μg RNA were used for cDNA synthesis and mixed with 1 μL 10 mM dNTPs, 1 μL of oligo(dT) primer, and filled up with water to 13 µL. The reaction was incubated at 65°C for 5 min and followed by 1 min on ice. 4 μL 5x First-Strand Buffer, 1 μL 0.1 M DTT and 1 μL SuperScriptTM III RT were added. After 60 min at 50°C, the reaction was inactivated for 15 min at 70°C. Isolated cDNA was stored at -20°C.

### DNA/expression constructs

Gephyrin Δ199-233 was generated by overlap-extension PCR (OEP) using subcloned isolated cDNA from Δ/Δ mice and gephyrin P1 wt in pQE80L48 as templates. The PCR product (primers: 5’ CCTCATGCCATTGACCTTTTACG 3’ and 5’ ATCATTAGTGCCTCAGAGCTCAA 3’) from the cDNA was subcloned into pJET2.1 following the CloneJET PCR Cloning Kit (Thermo Fisher Scientific). For OEP, three fragments (aa1-162, aa153-243, aa234-736) generated and fused by PCR by combination of the following primer pairs: 5’CCAT CACGGATCCGCATGCGAGCTCGGTACCCCAATGGCGACCGAGGGAA3’ and 5’GAA TTCTGCAGTCGACGGTACCGCGGGCCCATGGCGACCGAGGGAATGATCCTCA3’; 5’GAATGCTTTCAGTTCATACTGCCAGCTCTA3’ and 5’TTTGCAGCAAGAGCTGCTGCAGTTATAT3’; 5’AGCTAATTAAGCTTGGCTGCAGGTCGACCCTCATAGCCGTCCGATGACCA3’ and 5’ATGATCAGTTATCTAGATCCGGTGGATCCCTCATAGCCGTCCGATGACCATGACGT3’; 5’CCATCACGGATCCGCATGCGAGCTCGGTACCCCAATGGCGACCGAGGGAA3’ and 5’ATGATCAGTTATCTAGATCCGGTGGATCCCTCATAGCCGTCCGATGACCATGACGT3’. The final, fused fragment was cloned into pQE80L by Gibson assembly using SmaI. Gephyrin P1 wt and Δ199-233 from pQE80L constructs were cloned by Gibson assembly into pAAV_hSyn_mScarlet (a gift from Karl Deisseroth, Addgene plasmid #131001) using EcoRI. Gephyrin was cloned using PCR and XhoI and KpnI into pEGFP-C2 (Clontech).

Furthermore, DLC1 was cloned into pmCherry-C3 (Clontech) by PCR and ligation using restriction sites XhoI and HindIII.

For virus production the plasmids pAdDeltaF6 (a gift from James M. Wilson, Addgene plasmid #112867; http://n2t.net/addgene:112867; RRID: Addgene_112867) and pAAV2/1 (a gift from James M. Wilson, Addgene plasmid #112862; http://n2t.net/addgene:112862; RRID: Addgene_112862) were used. The expression constructs of LSβ in pQE70 and GlyRβICD in ptYB were described previously^2,34^.

### Purification of recombinant gephyrin, LSβ and βICD peptide

Gephyrin P1 wt and Δ199-233 in pQE80L were transformed into *E. coli* BL21. Bacterial cultures grew o/n at 37°C under shaking at 110 rpm. At an OD600 of approx. 0.3., expression was induced by supplementation with 250 nM IPTG. Cells were harvested by centrifugation for 10 min at 5 000 rpm (JLA 8.1 rotor) at 4°C after incubation for 21 h at 18°C and 90 rpm. Cell pellets were resuspended in 100 mL ME lysis buffer (300 mM NaCl, 50 mM Tris/HCl, 5 mM β-Mercaptoethanol, 0.05% Tween20, pH 7.4) with the addition of 1 tablet of EDTA-free protease inhibitor (Roche) and a spatula of lysozyme and stored at -80°C until use. Thawed pellets were lysed by alternating twice sonification (3 min at 4°C with 40% amplitude, 30 sec pulse, and 30 sec pause) and pressure lysis (1 000 to 1 500 bar, EmulsiFlex). All subsequent steps were performed at 4°C. The homogenate was centrifuged for 1 h at 16 000 rpm (JLA 16.250 rotor). The supernatant was applied twice on 20 mL Ni-NTA beads (pre-equilibrated with ME lysis buffer). The column was washed three times each, with 5 column volumes (CV) ME lysis buffer, then with 5 CV ME wash buffer 1 (300 mM NaCl, 50 mM Tris/HCl, 5 mM β Mercaptoethanol, 0.05% Tween20, 20 mM Imidazole, pH 7.4), and lastly with 5 CV ME wash buffer 2 (300 mM NaCl, 50 mM Tris/HCl, 5 mM β Mercaptoethanol, 0.05% Tween20, 40 mM Imidazole, pH 7.4). Elution was performed with 2 CV ME elution buffer (300 mM NaCl, 50 mM Tris/HCl, 5 mM β-Mercaptoethanol, 0.05% Tween20, 300 mM Imidazole, pH 7.4). The eluate was concentrated with an Amicon filter (50 kDa cut off) at 3 220 × g and pre-cleared for gel filtration at 17 000 × g. Preparative gel filtration was performed with an ÄKTApurifier (SuperdexTM 200 or SuperdexTM 200 HiLoad 16/60 prep grade, GE Healthcare) using SEC buffer (300 mM NaCl, 50 mM Tris/HCl, 5 mM β Mercaptoethanol, 5% glycerol, pH 7.4). Fractions of gephyrin trimers and oligomers were collected and concentrated by Amicon filters (50 kDa cut off) at 3 220 × g. Final proteins were snap frozen with liquid nitrogen, and stored at -80°C.

*E. coli* BL21 Rosetta star was used for the expression of His-LSβICD. The expression was performed as previously described34. Proteins were stored in 300 mM NaCl, 50 mM Tris/HCl, pH 7.4.

For the βICD peptide *E. coli* ER2566 (New England Biolabs) was used. Cultures grew at 37°C and 110 rpm until the OD600 reached 0.3. Induction was induced by 250 µM IPTG and expression performed for 20 h at 25°C and 110 rpm. Cells were harvested and stored at -80°C in ICD lysis buffer (300 mM NaCl, 50 mM Tris/HCl, 1 mM EDTA, 1x Protease Inhibitor (Roche), pH 7.4). Lysis of the thawed cells was performed by alternation of ultrasound (3 min at 4°C, 30 sec pulse, 30 sec pulse, 40% amplitude) and pressure lysis (EmulsiFlex, 1 000-1 500 bar, 4°C) twice. After centrifugation for 1 h at 16 000 × g (JLA 16.25 rotor) at 4°C, the supernatant was applied on a pre-equilibrated chitin matrix (New England Biolabs) and incubated 1.5 h at 4°C under shaking. The column was washed with 15 CV ICD buffer before 2.5 CV cleavage buffer (300 mM NaCl, 50 mM Tris/HCl, 1 mM EDTA, 50 mM DTT, pH 7.5) was applied an incubated for minimum 24 h at rt. The eluate was collected and another 2.5 CV cleavage buffer used for a second elution. The combined eluate was concentrated and filtered by Amicon filters with a cut-off of 10 kDa and 3 kDa. The buffer was exchanged to ITC buffer (300 mM NaCl, 50 mM Tris/HCl, pH 7.4) by dialysis over night at 4°C. The purified protein was stored at -80°C.

### Circular dichroism (CD) spectroscopy

Recombinant gephyrin was buffer exchanged to CD buffer (10 mM K_2_HPO_4_/KH_2_PO_4_, pH 7.5) and centrifuged for 10 min at 17 000 × g at 4°C. The concentration was adjusted to 0.2 mg/mL. Measurements were obtained using the JASCO J-715 CD Spectropolarimeter and data evaluated using Spectra Manager Version 2.

### Co-sedimentation assay

The previously described sedimentation assay^33^ was adapted as followed. Recombinant gephyrin and LSβ were pre-cleared by centrifugation at 17 000 × g for 3 min at rt. Gephyrin and LSβ were mixed 10:10 μM in a final volume of 50 μL of assay buffer (150 mM NaCl and 50 mM Tris/HCl, 5 mM TCEP, pH 7.5). After incubation of 10 min at rt, samples were centrifuged 10 min and 17 000 × g. The supernatant was added into a new tube and supplemented with 15 μL 5x SDS sample buffer. The pellet was supplemented with 50 μL assay buffer and 15 μL 5x SDS sample buffer.

### Isothermal titration calorimetry (ITC)

Gephyrin wt and Δ199-233 were buffer exchanged to ITC buffer (250 mM NaCl, 20 mM Tris/HCl, 5 mM β-mercaptoethanol, 5% glycerol, pH 7.5). Samples were pre-cleared by centrifugation for 10 min at 17 000 × g and the concentrations adjusted to 300 µM (βICD) and 30 µM (gephyrin). The ITC runs were performed with the MicroCal Auto-iTC200 (Malvern Panalytical) at 37°C with 40 individual injections (first injection: 0.5 µL in 1 sec; following injections: 1.25 µL in 2.5 sec), spacing in between injections of 2 min, and stirring speed of 750 rpm. βICD was titrated into gephyrin. Thermodynamic parameters were determined using the software Microcal analysis Origin 7.

### Analytical size exclusion chromatography (SEC)

Typically, 1 nmol of recombinant protein was supplemented with SEC buffer (300 mM NaCl, 50 mM Tris/HCl, 5 mM β-Mercaptoethanol, 5% glycerol, pH 7.4) in a final volume of 30 μL. Samples were centrifuged for 10 min at 17 000 × g and 4°C before injection. Runs were performed with Superose 6 Increase 5/150 GL in SEC buffer. The calibration curve was obtained using a protein standard containing thyroglobulin (669 kDa), ferritin (440 kDa), aldolose (158 kDa), conalbumin (75 kDa), and ovalbumin (44 kDa).

### Expression of EGFP-gephyrin and mCherry-DLC1 in COS7 cells

Coverslips were placed into 12-well plates and coated with collagen (0.2 mg/mL in DPBS) for 4 h at 37°C and afterwards washed with PBS twice. 40 000 COS7 cells (cultured in DMEM (+/+; Panexin Mix/Gln)) were seeded per well. Cells grew o/n at 37°C and were subsequently transfected using PEI with EGFP-gephyrin (pEGFPC2) and mCherry-DLC1 (pmCherry-C3). On the next day, the medium was replaced with fresh medium and expression was performed for another day (in total 48 h). Afterwards, cells were used for ICC.

### Culture of hippocampal neurons and expression of recombinant gephyrin

At E17.5, dissociated primary hippocampal cultures were prepared from C57BL/6NRj embryos. 90 000 cells were seeded on Poly-L-lysine coated 13 mm cover slips in 24-well plates. Neurons grew in neurobasal medium supplemented with B-27, N-2, and L-glutamine (Thermo Fisher Scientific). At 10 days *in vitro* (DIV), cells were transduced with 2.5 × 108 viral genome copies (GC) by exchanging a third of the medium to virus-containing medium. The virus was prepared and detected according to previous protocols [33]. At DIV14 the cells were either fixed or at DIV13-15 used for electrophysiology experiments. In case of electrophysiology measurements, 50 mM NaCl were added at DIV11 to maintain osmolarity. If cells should be used for Western blot analysis, cells were seeded Poly-L-lysine 24-well plates without cover slips. At DIV14, the medium was exchanged to 100 µL RIPA buffer (50 mM Tris/HCl, pH 7.5, 150 mM NaCl, 0.1% SDS, 1% IGEPAL CA360, 0.5% Sodium deoxycholate, 5 mM EDTA, 1x protease inhibitor (Roche)) and incubated for 10 min at rt under mild shaking. The detached cells were lysed by sonification (10 sec, 30%, 4°C) and centrifuged for 10 min at 4 000 × g. Approx. 30 µg of protein in the supernatant was used for SDS-PAGE.

### Immunocytochemistry (ICC)

Fixation was performed by replacing culture medium by 4% PFA in PBS. After incubation for 20 min at rt, PFA was removed and cells washed twice with 50 mM NH_4_Cl in PBS, each time 10 min at rt. Subsequently, cells were blocked for 1 h at rt with blocking solution (10% goat serum, 1% BSA, 0.2% TritonX100, in PBS). After washing cells with PBS for 5 min at rt, antibody incubation was performed for 1 h at rt. Antibodies were diluted in PBS. Afterwards, three washing steps with PBS were performed, each 10 min at rt. Treatment with secondary antibody and subsequent washing was performed in the same manner. In case of COS7 cells, staining with 300 nM DAPI in PBS was performed for 5 min at rt and an additional washing step of 10 min with PBS. Lastly, coverslips were mounted with Mowiol/Dabco and dried over night at rt. We used the following antibodies: rabbit anti-vesicular GABA transporter (vGAT) (1:1 000, #131003, SYSY), guinea-pig anti-GABA_A_Rγ2 (1:500, #224004, SYSY), rabbit anti-PSD95 (1:500; #ab18258; Abcam), guinea-pig anti-vesicular glutamate transporter (vGLUT) (1:1 000; #AB5905; Millipore), goat anti-rabbit AlexaFluor 647 (1:500, #A-21245, Invitrogen), goat anti-rabbit AlexaFluor 488 (1:500, #A-11075, Invitrogen), goat anti-guinea-pig AlexaFlour 647 (1:500; #ab150187; Abcam).

### Confocal microscopy and image analysis

Image stacks [0.33 μm z-step size, 10 stacks (=3 µm), 2048 × 2048 (144.77 × 144.77 μm)] were acquired on a Leica TCS SP8 LIGHTNING upright confocal microscope with an HC PL APO CS2 63×/1.30 glycerol objective, equipped with hybrid detectors (Leica HyD) and the following diode lasers: 405, 488, 552, and 638 nm. LIGHTNING adaptive deconvolution using “Mowiol” setting was applied. This form of adaptive image reconstruction is capable of theoretical resolutions down to 120 nm (lateral) and 200 nm (axial). Images were segmented and analyzed in an automated fashion using ImageJ/FIJI 1.53t. Therefore, the macros described by Liebsch et al.^32^ were used as a blueprint and adapted. Furthermore, we used the plugin JaCoP to determine Mander’s co-localization coefficients (MCCs).

### Electrophysiology experiments

Whole-cell patch-clamp recordings under current clamp and voltage clamp were conducted on primary cultures of murine hippocampal mScarlet- and Cre-moxBFP-tagged neurons between DIV13 and DIV15 at 30°C. Neurons were visualized with a fixed-stage upright microscope (Axio Examiner.D1, Carl Zeiss GmbH) using a water-immersion objective (Plan-Apochromat, 40×, 1 numerical aperture, 2.5 mm working distance, Carl Zeiss GmbH). The microscope was equipped with fluorescence and infrared differential interference contrast optics [50]. Red fluorescing gephyrin-positive cells were visualized with an X-Cite 120 illumination system (EXFO Photonic Solutions). Recordings were performed with an EPC9 patch-clamp amplifier (HEKA) controlled by the program PatchMaster (version 2 × 90.5, HEKA running on Windows 10). Data were recorded using a Micro1401 data acquisition interface (Cambridge Electronic Design Limited) and Spike 2 (version 7.08, Cambridge Electronic Design Limited). Data were sampled at 20 kHz and low-pass filtered at 10 kHz with a four-pole Bessel filter. Offline, the signal was smoothed by averaging the values in a ±0.001 s interval around any given datapoint. Electrodes with tip resistances between 4 and 6 MΩ were made from borosilicate glass (0.86 mm inner diameter; 1.5 mm outer diameter; GB150-8P, Science Products) with a vertical pipette puller (PP-830, Narishige).

During the recording, neurons were continuously superfused at a flow rate of ∼2.5 ml min−1 with a carbonated (95% O_2_ and 5% CO_2_) extracellular saline containing (in mM): 125 NaCl, 21 NaHCO_3_, 2.5 KCl, 2 MgCl_2_, 1.2 NaH_2_PO_4_, 10 HEPES, 5 glucose, adjusted to pH 7.2 with NaOH. For the voltage clamp recordings, glutamatergic input was blocked with 5 × 10−5 m DL-2-amino-5-phosphonopentanoic acid (DL-AP5; BN0086, Biotrend Chemikalien GmbH) and 10−5 M 6-cyano-7-nitroquinoxaline-2,3-dione (CNQX; C127, Sigma-Aldrich). Na^+^ mediated action potentials were blocked by 10−7 M tetrodotoxin (TTX; T-550, Alomone, Jerusalem BioPark). Current-clamp recordings were performed with a pipette solution containing (in mM): 140 K-gluconate, 10 KCl, 2 MgCl_2_, 10 HEPES, 0.1 EGTA, adjusted to pH 7.2 with KOH.

For voltage clamp recordings of miniature IPSCs (mIPSCs), the patch pipette solution contained (in mM): 133 CsCl, 2 MgCl_2_, 1 CaCl_2_, 10 HEPES, 10 EGTA, adjusted with CsOH to pH 7.2. The liquid junction potential between intracellular and extracellular solution for current-clamp (14.6 mV) and voltage-clamp (6.3 mV) recordings were calculated using the LJPcalc software (https://swharden.com/LJPcalc; arXiv:1403.3640v2; https://doi.org/10.48550/arXiv.1403.3640) and compensated accordingly.

Neurons were voltage clamped at −70 mV. Because of the high intracellular chloride concentration, GABA_A_ R-mediated currents were detected as inward currents.

Data were analyzed at 50 s intervals after the recording stabilized (∼10 min after obtaining whole-cell configuration). Data analysis was performed with Spike 2 (version 7.08, Cambridge Electronic Design Limited), Igor Pro 6 (version 6.37, Wavemetrics), and GraphPad Prism (version 10.2.3, GraphPad Software Inc.). mIPSC frequency, amplitude, and decay were determined offline. Events in voltage clamp traces were automatically detected by Easy Electrophysiology (developer pre-release version: v2.7.0, Joseph J Ziminski, www.easyelectrophysiology.com), when the signal crossed a threshold that was set and adjusted manually depending on the noise level of the signal. Events were classified as mIPSCs by template matching (correlation cutoff: 0.80; decay search period: 50 ms; baseline search period: 5 ms). The recording-specific template was calculated by fitting the average of prior threshold-detected and visually reviewed events. Parameters including frequency, amplitude, rise, and decay were determined. Statistical analysis between the genotypes was performed using a two-sided permutation test.

### SDS-PAGE, Coomassie and Western Blot

Samples were supplemented with 1-2.5x sample buffer (5 x: 250 mM Tris/HCl pH 6.8, 30% glycerol, 0.1% bromophenol-blue, 5% β-mercaptoethanol, 10% SDS) and incubated for 5 min at 95-98°C. SDS PAGE was performed with running buffer (25 mM Tris/HCl, 190 mM glycine, 0.1% SDS). As markers, the PageRulerTM Plus Prestained Protein Ladder or the unstained Protein Molecular Weight Marker were used.

For Coomassie staining, the SDS gel was stained with Coomassie stainer (30% EtOH, 10% acetic acid, 0.25% Coomassie brilliant blue R250) and destained first in destainer solution (30% EtOH, 10% acetic acid) and afterwards with water. Gel pictures were obtained using the ChemiDocTM MP Imaging System by BioRad.

For Western blot analysis, SDS gels were blotted semi-dry on PVDF membranes using transfer buffer (25 mM Tris/HCl, 320 mM glycine, 10% MeOH, pH 8.8) o/n at rt (25 mA). Blocking was performed for 1 h in blocking solution (10% milk powder in TBST buffer (50 mM Tris/HCl, 150 mM NaCl, 0.05% Tween20, pH 7.4)). Incubation in primary (antibody diluted in antibody solution (1% milk powder in TBST buffer)) was done for 1 h at rt. After washing three times for 5 min with TBST, incubation with the secondary antibody (diluted in antibody solution) was done for 1 h at rt. Afterwards blots were washed twice for 5 min with TBST, and once for 5 min with water. The membrane was developed using ECL and the ChemiDocTM MP Imaging System (BioRad).

When subsequent antibodies were used, membranes were incubated twice for 10 min each with stripping buffer (15 g/L glycine, 1 g/L SDS, 10 mL/L Tween20, pH 2.2), PBS, and lastly TBST. Then, antibody staining was performed as described in the previous section.

The following antibodies were used: mouse anti-GephyrinE (3B11; 1:10, self-made); rabbit anti-GAPDH (1:1 000, #G9545, Sigma), rabbit anti-PSD95 (1:1 000, #ab18258, Abcam); goat anti-mouse HRP-coupled (1:10 000, #AP181P, Sigma), goat anti-rabbit HRP-coupled (1:10 000, #AP187P, Sigma).

### Statistics

Statistical tests and methods used to determine significance and error bars are described in each figure legend. Data points exceeding two-fold standard deviation were determined as outliers. Numbers of replicates (N) are indicated in each figure legend. Note, that in case of neuro cell culture experiments, 10 cells per condition (technical N=10) from 3 individually prepared pups (biological N=3) were analyzed (total N=30). In case of COS7 cells, 5 cells per condition (technical N=5) from 3 individual seedings (biological N=3) were analyzed (total N=15). For electrophysiology experiments (voltage clamp recordings), we used cell cultures from 3 individual pups (biological N=3) and analyzed 7-9 individual cells (technical N = 9 for wt and N = 7 for gephΔ199-233).

## Acknowledgements

Technical help from Julia Reich is greatly appreciated. We thank Simon Tröder and Branko Zevnik as well as the whole CECAD TCU for the generation of the mouse line. We also thank Franziska Neuser and the Center for Mouse Genetics (CMG) of the University of Cologne for the organization of mice experiments. Lastly, we thank the imaging facility of the Cologne Biocentre of the University of Cologne. This work was financed by the german research foundation (DFG) (RTG2550 reloc to JR).

## Conflict of interest

The authors declare no conflict of interest.

## Study approval

We complied with all ethical regulations for animal testing and research. Experiments were approved by the local research ethics committee (German, Landesamt für Natur, Umwelt und Verbraucherschutz (LANUV) Nordrhein-Westfalen, reference 2016.A466).

## Notes

### Competing Interest Statement

The authors have declared no competing interest.

