## Supplementary Data for "GephyrinΔ199-233 - an epileptogenic microdeletion"

### Supplementary figures

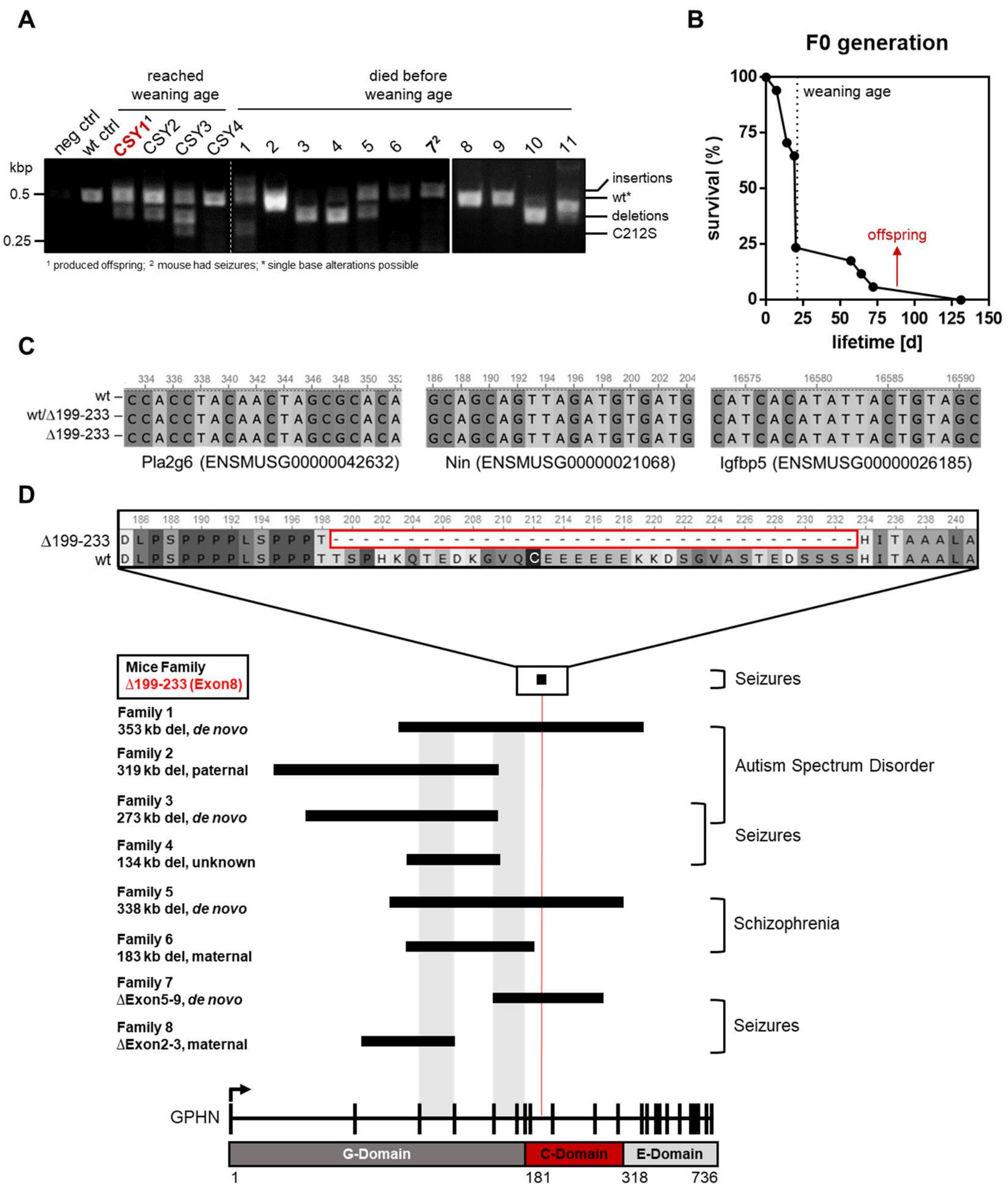


<u>Figure S1: Generation of gephyrin C212S and gephyrin Δ199-233 KI mice</u>

A) Genotyping of founder mice. PCR was designed to obtain amplicons around the area of the gRNA binding site. C212S point mutation identified, but several mice showed also insertions and deletions. Amplicons of expected wt-size can still carry single base alterations. Only mouse CSY1 was able to produce offspring. Mouse 7 had to be sacrificed after seizures. B) Survival rate [%] of the founder generation (F0). 25% mice reached weaning age. C) Alignment of off-target regions of the CRISPR/Cas guide RNA. Homozygous wt and Δ199-233 and heterozygous mice in comparison. D) Comparison of gephyrin deletion Δ199-233 in the new mice line with human patient deletions showing corresponding neuronal impairments. Deletions assigned to the different exons of the GPHN gene structure. Common exons shadowed in light grey. Specific region of gephyrin Δ199-233 enlarged.

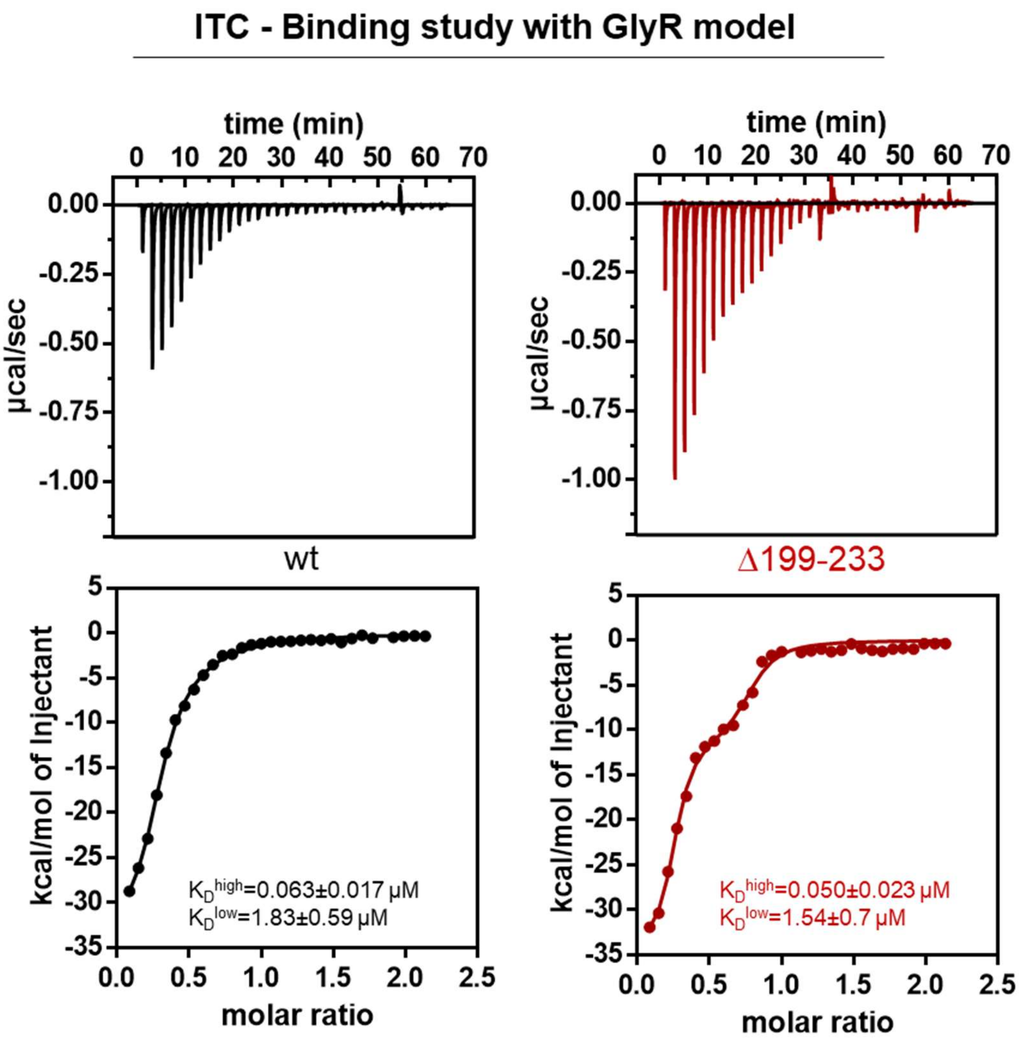


Figure S2: Receptor interaction study of gephyrin Δ199-233

A) Isothermal calorimetry titration experiment using GlyRβICD peptide as a model for the GlyR. The peptide was titrated into gephyrin. Data has been fitted with a two-binding site model giving a high and a low affinity dissociation constant ($K_D^{high}$; $K_D^{low}$).
